# Ryegrass mottle virus complete genome determination and development of infectious cDNA by combining two methods – 3′ RACE and 5′ RACE-seq

**DOI:** 10.1101/2023.06.04.543628

**Authors:** Ina Balke, Ivars Silamikelis, Ilze Radovica-Spalvina, Vilija Zeltina, Gunta Resevica, Davids Fridmanis, Andris Zeltins

**Author notes:** corresponding author (IB).

## Abstract

Sobemovirus ryegrass mottle virus (RGMoV) is a single-stranded positive virus with a 30 nm viral particle size. It exhibits *T=3* symmetry, with 180 coat protein (CP) subunits forming the virus structure. The RGMoV genome comprises five open reading frames, encoding P1, Px, a membrane-anchored 3C-like serine protease, a virus genome-linked protein, P16, an RNA-dependent RNA polymerase, and a coat protein. The RGMoV genome size varies, ranging from 4175 nt (MW411579.1) to 4253 nt (MW411579.1) in deposited sequences. An earlier deposited RGMoV complete genome sequence of 4212 nt length (EF091714.1) was utilized to develop an infectious complementary DNA (icDNA) construct for *in vitro* gRNA transcription from the T7 promoter. However, when the transcribed gRNA was introduced to oat plants, it failed to induce viral infection. This indicated the potential absence of certain sequences in either the 5’ or 3’ untranslated regions (UTR) or both. To resolve this, the complete sequence of the 3’ UTR was determined through 3’ end RACE, while the 5’ UTR was identified using high-throughput sequencing (HTS) - 5’ RACE-seq. Only the icDNA vector containing both newly identified UTR sequences proved infectious, resulting in classical viral infection symptoms and subsequent propagation of progeny viruses, exhibiting the ability to cause repeated infection in oat plants after at least one passage. The successful generation of the icDNA highlights the synergistic potential of utilizing both methods when one approach alone fails. Furthermore, this study demonstrates the reliability of HTS as a method for determining the complete genome sequence of viral genomes.

## Introduction

Ryegrass mottle virus (RGMoV) is classified under the genus *Sobemovirus* within the recently established family *Solemoviridae*, which also includes the genera *Polemovirus*, *Polerovirus*, and *Enamovirus* [1]. Genus *Sobemovirus* contain 21 assigned species and two related viruses. The *Sobemovirus* genus name was derived from the reference species ***<u>So</u>****uthern **<u>be</u>**an **<u>mo</u>**saic virus*, while the family name *Solemoviridae* combines the founding genera ***<u>So</u>****bemovirus* and *Po**<u>lemo</u>**viru* [2]. RGMoV was first isolated from stunted Italian ryegrass (*Lolium multiflorum*) and cocksfoot (*Dactylis glomerate*) showing symptoms of leaf mottling and necrosis [3]. The virus is composed of an isometric particle with a diameter of 28 nm and contains a single-stranded positive-sense genomic RNA (gRNA) weighing approximately 1.5×106 Da. The virion comprises a single coat protein (CP) with a molecular weight (MW) of 25.6 kDa [4]. The capsid structure exhibits *T=3* symmetry, consisting of 180 CP subunits arranged in a canonical jellyroll β-sandwich fold. Compared to other sobemoviruses, the RGMoV particle is 3% smaller in diameter, and the virion cavity is 7% smaller [5]. The gRNA of RGMoV has a virus-genome linked protein (VPg) at the 5’ terminus, while the 3’ end lacks a poly(A) tail [4]. The gRNA encodes five open reading frames (ORFs) [6, 7]. ORF1 codes for P1, a zinc finger protein with movement protein function and potential long-distance RNA silencing suppressor roles [8, 9]. ORFX is responsible for establishing the infection [7]. ORF2a encodes the polyprotein P2a, which consists of three protein domains: a membrane anchor 3C-like serine protease (Pro) with a typical chymotrypsin-like protease structure [10], a "natively unfolded" VPg protein [11] that interacts with eukaryotic translation initiation factor eIF(iso)4G [12, 13], serving for ribosome recruitment [13], and a protein with a calculated MW of 16.8 kDa [11], possessing nucleic acid binding properties and a novel Mg^2+^-dependent ATPase activity [14]. ORF2b, located in the -1 frame relative to ORF2a, encodes the RNA-dependent RNA polymerase (RdRp) [6]. *In vitro* studies have demonstrated that the RdRp of Sesbania mosaic virus (SeMV) initiates progeny RNA synthesis in a primer-independent replication manner. The presence of the ACAA motif at the 3’-end of the negative-sense strand suggests that it might function as a promoter or enhancer for replication [15-17]. ORF3 of RGMoV contains a coding sequence for the *CP*, which is translated from a subgenomic RNA (sgRNA) translated from a sgRNA [5, 18]. In the case of CfMV, the CP suppresses RNA silencing but is not required for systemic movement [19]. The translation of ORFx is predicted to rely on a non-AUG initiation mechanism, while the translation of ORF2a depends on a leaky scanning mechanism [7]. The processing of the polyprotein 2a by the Pro occurs *in cis* at E117/A118 [10]. However, it also occurs *in trans* at E315/S316 and E393/S394, resulting in the release of mature Pro, VPg, and P16 domains [10, 20].

It is believed that each virion of sobemoviruses encapsidates one gRNA molecule and one sgRNA molecule. Some sobemoviruses have been reported to encapsidate viroid-like satellite RNAs (satRNAs), which are dependent on a helper virus for replication. This phenomenon has been observed in several sobemoviruses, including lucerne transient streak virus (LTSV), rice yellow mottle virus (RYMV), subterranean clover mottle virus (SCMoV), Solanum nodiflorum mottle virus (SNMoV), and velvet tobacco mottle virus (VTMoV). Cocksfoot mottle virus (CfMV) has been shown to encapsidate the satRNA of LTSV during infection in two monocotyledonous species, *Triticum aestivum* and *Dactylis glomerate* [21]. However, for CfMV, only the presence of defective interfering RNA (diRNA) has been detected encapsidated in viral particles [22].

The genome size of RGMoV deposited in GenBank, labeled as "complete genome," ranges from 4175 nt (GenBank ID MW411579.1) to 4253 nt (GenBank ID MW411579.1). The distribution of RGMoV is no longer limited to Japan. Through high throughput sequencing (HTS), RGMoV has been detected in various samples worldwide, including the United Kingdom (GenBank ID MW588198.1), USA (GenBank ID OK424596.1), Australia (GenBank ID MT129760.1), and Ecuador (GenBank ID MW411579.1; [23]). This widespread presence of RGMoV in different regions marks it as a plant pathogen of concern. The identification of RGMoV in these diverse locations suggests the necessity for ongoing monitoring of this virus, especially considering that several other sobemoviruses are of high economic importance, such as rice yellow mottle virus (RYMV) and papaya lethal yellowing virus [24].

The genome of RGMoV has undergone revision in previous studies, specifically concerning the amino acid (aa) sequence of the Pro [4, 6]. The coding region of RGMoV has been re-verified through re-sequencing efforts [6, 25]. However, the concern remains regarding the 5’ and 3’ untranslated regions (UTR) of the genome. Currently, there are no reports on infectious complementary viral DNA (icDNA) for RGMoV, making the true complete genome sequence unknown. While RGMoV shares some physical and biological properties with CfMV, it is serologically distinct from CfMV, as well as other viruses such as cocksfoot mild mosaic virus, Cynosurus mottle virus, and Phleum mottle virus found in European countries [4]. Despite these serological differences, RGMoV shares general properties with other sobemoviruses that infect grasses [3, 26]. The transmission vector for RGMoV is still unknown, but like other sobemoviruses [24], it can be transmitted through the mechanical wounding [5].

The development of the first icDNA of brome mosaic virus (BMV) by Ahlquist et al. in 1984 paved the way for the establishment of multiple reverse genomics systems based on plant viruses [27]. The construction of infectious clones of plant viruses has played a crucial role in advancing our understanding of these viruses at the molecular level. It has enabled the characterization of viral replication, host range, movement, pathogenicity, and the functional roles of different genomic regions. Moreover, infectious clones have been utilized as gene expression vectors and tools for studying virus-induced RNA silencing. They have also contributed to the identification of virus-host interaction pathways [28]. In the case of sobemoviruses, icDNA has been successfully generated for only four viruses: RYMV [29], CfMV [30], Sowbane mosaic virus (SoMV; previous known also as Rubus chlorotic mottle virus) [31], and SeMV [16]. These icDNA constructs have provided valuable tools for studying the biology of these viruses and investigating their interactions with host plants.

We present the comprehensive genome sequence of RGMoV and describe the successful generation of the icDNA for this virus. The complete RGMoV genome consists of a gRNA with a length of 4287 nucleotides (nt). The gRNA contains a 117 nt long 5’ UTR and a 261 nt long 3’ UTR. Furthermore, we successfully reinfected plants with purified virus obtained from the icDNA capped transcript. The infection was observed after at least one passage, indicating the viability and infectivity of the icDNA-derived RGMoV. This achievement provides valuable insights into the biology and pathogenicity of RGMoV and opens up possibilities for further investigations into the virus-host interactions and molecular mechanisms underlying RGMoV infections.

## Materials and Methods

### Virus propagation and extraction

RGMoV (National Institute of Agrobiological Sciences, Japan, MAFF No. 307043) was maintained and propagated in oats (*Avena sativa* cv. Jaak), which were grown in a Climatic Test and Plant Growth Chamber MLR-351H (Sanyo, Osaka, Japan) under controlled conditions. The oats were grown at temperatures ranging from 19 to 23 °C with a 17-hour light period.

To subculture the virus, sap inoculation was performed on *A. sativa* plants using extracts obtained from RGMoV-infected plant leaves. The infected leaves were stored at -80 °C and ground in 15 mM potassium phosphate buffer (pH 7.4) mixed with diatomaceous earth. The mixture was gently rubbed on the upper leaf surface of 2-week-old oats, targeting the second, third, and fourth leaves.

RGMoV was purified from oat plants displaying mottling and necrotic symptoms on their leaves [3], following a previously described protocol with some modifications as described earlier [5]. In brief, the virus was extracted from 12.82 g of oat plants, which were homogenized in 5 volumes of 0.075 M potassium phosphate buffer (pH 5.5). The sap was filtered to remove oat debris, and chloroform was added (0.4 volumes) to the filtrate. The mixture was gently agitated at room temperature (RT) for 30 min, followed by centrifugation at 5000 rpm (3214 × g) for 10 min at 4 °C. The upper phase was collected and transferred to new tubes, and 25% ammonium sulfate was added to achieve a final concentration. The solution was mixed ON at 4 °C. The ammonium sulfate concentration was then increased to 50% (final concentration) and incubated with agitation for 3 h. Subsequently, the solution was centrifuged at 11,000 rpm (15,557 × g) for 20 min at 4 °C. The resulting pellet was dissolved in 15 mM potassium phosphate buffer (pH 5.5) and dialyzed ON at 4 °C in the same buffer.

The virus was further purified using sucrose density gradient centrifugation, following a similar protocol to the purification of virus-like particles from RYMV and CfMV [32]. Fractions containing the virus were dialyzed against 200 volumes of 15 mM potassium phosphate buffer (pH 5.5) and concentrated through ultracentrifugation at 72,000 rpm (280,000 × g) for 1 h at 4 °C. The purified virus particles were solubilized in 15 mM phosphate buffer (pH 5.5), analyzed by transmission electron microscopy (TEM), and stored at 4 °C for further studies.

### Genomic and total RNA extraction for 5’ and 3’ RACE, HTS and RT-PCR

For the determination of the 5’ and 3’ UTRs of the RGMoV genome using 5’ and 3’ RACE (rapid amplification of cDNA ends), the viral gRNA was isolated from 300 µg of purified viral particles. The viral particles were disassembled in a dissociation solution containing 100 mM NaCl, 10 mM TRIS-HCl (pH 8.0), 0.5% SDS, 1 mM EDTA, and 40 µg of Proteinase K (Thermo Fisher Scientific, Waltham, MA, USA). The mixture was incubated at 37 °C for 1 h.

RNA extraction was performed by adding one volume of phenol-chloroform solution (1:1) to the disassembled viral particles, followed by centrifugation at 13,200 rpm (16,100 × g) for 10 min. The RNA was then precipitated by adding 1/10 volume of potassium acetate (pH 5.3) and two volumes of 96% ethanol. The mixture was incubated at -20 °C for 30 min and centrifuged at 13,200 rpm (16,100 × g) for 10 min. The RNA pellet was washed with one volume of 75% ethanol and centrifuged at 13,200 rpm (16,100 × g) for 5 min. The RNA pellet was air-dried for 10 min at RT and dissolved in 30 µl of DEPC-treated water (Thermo Fisher Scientific, Waltham, MA, USA). The concentration of the extracted RNA was measured using a NanoDrop-1000 spectrophotometer (Thermo Fisher Scientific, Waltham, MA, USA) and stored at -80 °C until further use.

For high-throughput sequencing (HTS), viral gRNA was purified from a solution of viral particles with a concentration of 1 mg/ml. The viral particles were treated with Benzonase (25 units/µl; Novagen, Bad Soden, Germany) to remove nucleic acid from the outer space of the viral particles and the solution. The viral gRNA was then extracted from the purified viral particles using TRI REAGENT® according to the provided protocol. The purified RNA was dissolved in 30 µl of RNase-free water. The concentration of the purified RNA was measured using both NanoDrop-1000 and Qubit 2.0 (Thermo Fisher Scientific, Waltham, MA, USA) with the Qubit RNA high sensitivity (HS) assay kit (Thermo Fisher Scientific, Waltham, MA, USA). The RNA was stored at -80 °C until further use.

Total RNA for reverse transcription polymerase chain reaction (RT-PCR) was isolated from 100 mg of oat leaves using TRI REAGENT® (Sigma-Aldrich, St. Louis, MO, USA) according to the provided protocol. The purified RNA was dissolved in 30 µl of DEPC-treated water. The concentration of the total RNA was measured using NanoDrop-1000 and analyzed on a 0.8% native agarose gel (NAG). The total RNA was stored at -80 °C.

### RGMoV gRNA 5’ and 3’ RACE verification by Sangers sequencing

To determine the complete genome sequence of RGMoV and develop the icDNA construct, the 3’ UTR of the virus were determined using the SMARTer®RACE 5’/3’ Kit (Takara Bio, Kusatsu, Japan). 2.4 µg of RGMoV gRNA was polyadenylated using Poly(A)polymerase (600 u/µl; USB, Cleveland, OH, USA). The polyadenylated RGMoV gRNA was used for first-strand cDNA synthesis according to the manufacturer’s protocol of the SMARTer®RACE 5’/3’ Kit. The 5’ and 3’ ends of the RGMoV genome were amplified separately. The 5’ end was amplified using the genome-specific (GenBank Accession number EF091714.1 [6]) primer RG-P1-R (5’ GGATCCATGATGTCTAGTCCAAGACTGCCCT 3’) and the kit-provided 10x Universal Primer Mix (10xUPS). The 3’ end was amplified using the genome-specific primer RG-seqCP-F (5’ ATCTGGGCAAGGGTCCCGCTATTCGAAG 3’) and 10xUPS. The PCR products of the 5’ and 3’ ends were extracted from the gel after analysis in a 0.8% native agarose gel using the GeneJET Gel Extraction Kit (Thermo Fisher Scientific, Waltham, MA, USA). Adenine overlaps were added to the PCR products using Taq polymerase (Thermo Fisher Scientific, Waltham, MA, USA), and the products were cloned into the linearized vector pTZ-57 using the InsTAclon PCR Cloning Kit (Thermo Fisher Scientific, Waltham, MA, USA). This step resulted in the construction of plasmids named pTZ-5’end and pTZ-3’end. The plasmid constructs pTZ-5’end and pTZ-3’end, containing the PCR fragments, were sequenced using the Sanger sequencing method. The sequencing reactions were performed using the ABI PRISM BigDye Terminator v3.1 Ready Reaction Cycle Sequencing Kitt (Thermo Fisher Scientific, Waltham, MA, USA), and the sequences were determined using an ABI PRISM 3130xl sequencer (Thermo Fisher Scientific, Waltham, MA, USA). The corresponding primers M13seq-F (5’ GCC AGG GTT TTC CCA GTC ACG A 3’) and M13seq-R (5’ GAG CGG ATA ACA ATT TCA CAC AGG 3’) were used for sequencing. The obtained sequencing reads were assembled using SeqMan software (DNASTAR, Madison, WI, USA) to generate the complete genome sequence of RGMoV.

### RGMoV gRNA RNA-seq next-generation sequencing library preparation for HTS on the Ion Torrent Personal Genome Machine™ (PGM™) Sequencer

Before HTS library preparation, purified gRNA from benzonase-treated virus material was treated with DNaseI (Thermo Fisher Scientific, Waltham, MA, USA), and analyzed using an Agilent 2100 bioanalyzer (Agilent Technologies, Santa Clara, CA, USA) with an Agilent HS RNA Kit (Agilent Technologies, Santa Clara, CA, USA) to assess the level of fragmentation. The gRNA was fragmented into approximately 200 nt long fragments using an S220 Focused-ultrasonicator (Duty Cycle 10%; Intensity 5; Peak Incident Power 175 Watts; Cycles per Burst 200; Processing time 200 s; Covaris, Woburn, MA, USA). The fragmented RNA was analyzed on an Agilent 2100 bioanalyzer with an Agilent RNA 6000 Pico kit (Agilent Technologies, Santa Clara, CA, USA) to determine the length distribution of the fragments. The concentration of the fragmented RNA was measured using a Qubit 2.0 with a Qubit RNA HS assay kit. Hybridization of the fragmented RNA (100 ng) was performed with a barcode from the Xpress RNA-Seq Barcode 1-16 Kit (Thermo Fisher Scientific, Waltham, MA, USA), Ion Adaptor Mix, and Hybridization Solution from the Ion Total RNA-Seq Kit (Thermo Fisher Scientific, Waltham, MA, USA) according to provided protocol. The hybridization reaction took place in a Veriti 96-Well Thermal Cycler (Thermo Fisher Scientific, Waltham, MA, USA) at 65 °C for 10 min and 30 °C for 5 min. Ligation was then carried out according to the Ion Total RNA-Seq Kit protocol. RT-PCR was performed at 42 °C for 30 min in a Veriti 96-Well Thermal Cycler using the Ion Total RNA-Seq Kit protocol. The cDNA library was purified, and size selection was performed using the Agencourt AMPure XP reagent (Thermo Fisher Scientific, Waltham, MA, USA) as instructed in the protocol. The purified cDNA library was amplified using AmpliTaq DNA Polymerase (Thermo Fisher Scientific, Waltham, MA, USA) and Ion 5’PCR and Ion 3’PCR primers on a Veriti 96-Well Thermal Cycler following the Ion Total RNA-Seq Kit protocol. The amplified DNA was then purified using the PureLink PCR Micro Kit (Thermo Fisher Scientific, Waltham, MA, USA) following the manufacturer’s protocol. The purified DNA library was analyzed using an Agilent 2100 bioanalyzer with an Agilent DNA 1000 Kit (Agilent Technologies, Santa Clara, CA, USA) and a Qubit 2.0 with Qubit 1X dsDNA HS Assay Kits (Thermo Fisher Scientific, Waltham, MA, USA) to determine the concentration of the library.

Emulsion PCR was prepared using the Ion OneTouch System (Thermo Fisher Scientific, Waltham, MA, USA) according to the Ion OneTouch 200 Template Kit v2 DL protocol (Thermo Fisher Scientific, Waltham, MA, USA). Sequencing was performed on the Ion 314 Chip V2 (Thermo Fisher Scientific, Waltham, MA, USA) using the Ion PGM Sequencer (Thermo Fisher Scientific, Waltham, MA, USA) and Ion PGM Sequencing 200 Kit (Thermo Fisher Scientific, Waltham, MA, USA) following the standard protocol.

### HTS data analysis

After the gRNA RNA-seq library preparation, the sequencing reads underwent additional processing and analysis steps. Adapter, barcode and quality trimming was performed with cutadapt 4.2 [33] and fastp 0.23.2 [34]. Reads with length at least 50 bp reads were retained for further analyses. The trimmed reads were used for contig assembly using the SPAdes genome assembler v3.15.5 [35] The assembly was performed in "rnasviralSPAdes" mode with the "iontorrent" flag set on, indicating the specific sequencing platform used. Bowtie2 (version 2.3.5.1) [36] was used for read mapping against the assembled contigs and reference sequences. In order to identify reads related to additional sequences in the 5’ UTR region, manual sequence alignments were performed using SeqMan software. The resulting data and analyses were visualized using matplotlib 3.6.3 [37].

### RGMoV icDNA vector construction under *T7* polymerase promoter

The cDNA construct with RGMoV sequence corresponding to deposited in GenBank under Accession number EF091714.1 was reported earlier [6]. The 3’ UTR with an additional 52 nt sequence obtained by 3’ RACE was amplified from the pTZ-3’end plasmid. iProof High-Fidelity DNA Polymerase (500 U; Bio-Rad, Hercules, CA, USA) was used for the amplification, along with primers RG-seqCP-F and RG-3pN-Bst-R (5’ CAGTATACTACAACCCCTAGCGTGGAATGATCCTACCCTAGGCA 3’). The PCR reaction was performed on a Veriti 96-Well Thermal Cycler at an annealing temperature of 55 °C. The resulting PCR product was extracted from the gel using the GeneJET Gel Extraction Kit and adenine overlaps were added using Taq polymerase. The PCR product was then cloned into the linearized vector pTZ-57 using the InsTAclon PCR Cloning Kit, generating the plasmid construct pTZ-3’end-PCR. The clones containing the PCR fragment were verified by Sanger sequencing. The pTZ-3’end-PCR clone and a plasmid containing RGMoV gRNA were digested with BamHI restriction enzyme (Thermo Fisher Scientific, Waltham, MA, USA). The DNA fragments were extracted from the gel by GeneJET Gel Extraction Kit and ligated to generate a cDNA construct named pJET-RGMoV-cDNAp-3end. The constructed plasmid was analyzed by restriction analysis and verified by Sanger sequencing. Two variants of the 5’ end sequence were identified by HTS. The first construct was developed based on a 36 nt fragment (5UTR-new). A two-step PCR was performed to amplify the 5UTR-new fragment. The first PCR was carried out using primers RG-5UTR-new-F (containing *T7* promoter sequence; 5’ACCATCGCGATAATACGACTCACTATAGGG ACCTCTCTATGGGCA GTCTCCTCTCTATGGCAGTCGACAAA 3’) and RG-5UTR-new-R (5’TCACCGGATAACGGGTTCAATAGAGTTAATTTAATAACTCTATTTGTCGACTGCC ATAGAGAGGAGACTG 3’) and Pfu polymerase (Thermo Fisher Scientific, Waltham, MA, USA). The PCR product was extracted from the gel. The second PCR was amplified from the purified PCR product and a plasmid containing the RGMoV 1^st^ gRNA fragment, pTZ-1fr [6]. After 5 cycles of amplification with annealing temperature 55 °C primers RG-5UTR-new-F and RG2-P1-HindIII-R (5′ AAGCTTATGATGTCTAGTCCAAGACTGCCCTCGA 3′) were added followed by 25 cycle amplification at annealing temperature at 55 °C. added using the GeneJET Gel Extraction Kit, adenine overlaps were added, and it was cloned into the linearized vector pTZ-57 using InsTAclon PCR Cloning Kit. The resulting plasmid construct was named pTZ-5UTR-new and was verified by Sanger sequencing. The pTZ-RGMoV-5UTR-new and pJET-RGMoV-cDNA-3end constructs were cut with Eco91I (BstEII; Thermo Fisher Scientific, Waltham, MA, USA) and PstI (Thermo Fisher Scientific, Waltham, MA, USA) restriction enzymes. The DNA fragments were extracted from the gel by GeneJET Gel Extraction Kit and ligated to obtain the construct pTZ-T7-RGMoV-cDNA-3end-5UTR-new-PGM. A 5’ end construct with a 19 nt addition (5’UTR-new-short) was created using the pTZ-RGMoV-5UTR-new plasmid as a template. PCR amplification was performed with Pfu polymerase using primers RG-P1-R and RG-5UTR-new-short-F (containing the *T7* promoter sequence; 5’ ACCATCGCGATAATACGACTCACTATAGGG ACCTCTCTATGGGCAGT 3’). The PCR product was extracted from the gel, adenine overlaps were added, and it was cloned into the pTZ-57 cloning vector. The resulting plasmid, pTZ-RG-5UTR-new-short, was verified by Sanger sequencing. pTZ-RG-5UTR-new-short and pTZ-T7-RGMoV-cDNA-3end-5UTR-new-PGM were subjected to digestion with Mph1103I (NsiI; Thermo Fisher Scientific, Waltham, MA, USA) and Eco91I restriction enzymes. The resulting RG-5UTR-new-short fragment was then ligated into the pTZ-T7-RGMoV-cDNA-3end-5UTR-new-PGM plasmid. This ligation process generated the final icDNA construct named pTZ-T7-RGMoV-cDNA-3end-5UTR-new-short-PGM.

### RGMoV gRNA *in vitro* synthesis

Before RNA synthesis, all developed icDNA constructs, including pJET-RGMoV-cDNAp, pJET-RGMoV-cDNAp-5UTR-new-PGM, pJET-RGMoV-cDNAp-5UTR-new-short-PGM, pJET-RGMoV-cDNAp-3end, pTZ-T7-RGMoV-cDNA-3end-5UTR-new-short-PGM, and pTZ-T7-RGMoV-cDNA-3end-5UTR-new-PGM, were subjected to linearization by cutting with specific restriction enzymes. The plasmids (10 µg) pJET-RGMoV-cDNAp and pJET-RGMoV-cDNAp-5UTR-new-PGM were digested with HindIII (Thermo Fisher Scientific, Waltham, MA, USA), while pJET-RGMoV-cDNAp-3end, pTZ-T7-RGMoV-cDNA-3end-5UTR-new-short-PGM, and pTZ-T7-RGMoV-cDNA-3end-5UTR-new-PGM were digested with Bst1107I (Thermo Fisher Scientific, Waltham, MA, USA). To verify the success of the linearization process, the purified DNA was analyzed using native agarose gel electrophoresis (NAGE) in a 0.8% agarose gel stained with ethidium bromide. The concentration of the linearized DNA was estimated using a NanoDrop-1000 spectrophotometer.

Capped gRNA was synthesized using either the RiBoMax Large Scale RNA Production Systems (Promega, Madison, Wis, USA) carried out at 30 °C according to protocol or the TranscriptAid T7 High Yield Transcription Kit (Thermo Fisher Scientific, Waltham, MA, USA) following the respective protocol provided by the manufacturers for *in vitro* RNA transcripts with or without a cap. For the transcripts with a cap, the cap analogue used was Ribo m7G Cap Analog (40 mM; A254 units, Promega, Madison, Wis, USA, P171B). After synthesis, the RNA was purified using the RNeasy Plant Mini kit (Qiagen, Hilden, Germany), following provided protocol. To verify the integrity of the synthesized RNA, it was analyzed using NAGE in a 1% agarose gel stained with ethidium bromide. The concentration of the purified RNA was estimated using a NanoDrop-1000 spectrophotometer.

### RGMoV gRNA transcript coating on gold particles for Helios Gene Gun

To prepare the tubing for gene gun delivery one tubing and 25 mg of Au particles with 1 µm Ø was used from Helios Gene Gun Optimization Kit (Bio-Rad, Hercules, CA, USA). 10 µg of the synthesized RNA transcript was used four labeling Au particles. Tubing Prep Station (Bio-Rad, Hercules, CA, USA) was used for tubing coating with Au-RNA particles. All procedures were performed as described in the Helios Gene Gun System instruction manual.

### Oat inoculation with gRNA transcript

Oat plants were inoculated with the gRNA transcript in the first leaf at the second leaf stage using two different methods: mechanical rubbing and bombardment with Au particles coated with RNA. Control plants were mock-inoculated with DEPC-treated water.

The gRNA transcript, at a quantity of 2.5 µg, was mechanically rubbed onto the first leaf of oat plants using Carborundum (330 grit, Thermo Fisher Scientific, Waltham, MA, USA) as an abrasive agent. The Au-RNA particles were then used for bombardment onto the oat plants using the Helios Gene Gun (Bio-Rad, Hercules, CA, USA) at a pressure setting of Hpsi=200. The first inoculation was performed when the oat plants reached the second leaf stage. After one week, a second inoculation was conducted in the first leaf. Another week later, a third inoculation was performed at the third leaf stage in the second leaf. Oat plants were grown under the same conditions as those used for virus propagation. The plants were allowed to grow for four weeks after the third inoculation. Samples for SDS-PAGE, western blot (WB), and total RNA extraction for RT-PCR were collected from the fourth leaf of the oat plants. RT-PCR was conducted using the Verso 1-Step RT-PCR Hot-Start kit (Thermo Fisher Scientific, Waltham, MA, USA). The primers used for RT-PCR were RGCP-NdeI-F (5’ ATGGCAAGGAAGAAGGGCAAATCGGCCA 3’) and RGCP-Pich-EcoRI-R (5’ GGAATTCTCACTGGTTGATTGTGACATCAACCGGA 3’). The RT-PCR protocol provided by the kit manufacturer was followed for the amplification.

### Transmission electron microscopy (TEM)

Purified virus samples were adsorbed onto carbon formvar-coated grids (300 mesh Copper/Palladium 3.05 mm; Laborimpex, Forest, Belgium) or carbon formvar-coated grids (hexagonal 200 mesh nickel 3.05 mm; Laborimpex, Forest, Belgium) and negatively stained with a 0.5% uranyl acetate aqueous solution. The stained grids were examined using a JEM-1230 TEM (JEOL, Tokyo, Japan) at an accelerating voltage of 100 kV as described previously [38].

### Oat leaf extract analysis by SDS-PAGE and Western blot

100 mg of oat leaf was ground in a 1.5 ml tube with 200 µl 2xLaemmli sample buffer (100 mM Tris-HCI (pH 7.0), 2% SDS, 50% glycerol, 0.005% Bromophenol Blue, 2% mercaptoethanol) by micro pestle. Samples before loading on 12.5% dodecyl sulphate polyacrylamide gel [SDS-PAG; [32]] were incubated at 95 °C for 10 min. After SDS-PAG electrophoresis (SDS-PAGE) gel was stained with R250 (10% (v/v) ethanol, 10% (v/v) glacier acid, 0.1% (w/v) R250).

WB was performed on a semi-dry WB device transferred to the Amersham Protran 0.45 mm nitrocellulose blotting membrane (GE Healthcare) at 52 mA for 45 min. The membrane was blotted by a 1% alkali-soluble casein solution (Merck–Millipore) for 1 h at RT on Mixer 820 (Boule Medical, Stockholm, Sweden) at speed 60 Hz 18 rpm/8 rpm. After blocking the membrane was incubated ON at 4 °C in 1% alkali-soluble casein solutions with polyclonal anti-RGMoV (1:1000; produced in-house). The membrane was washed with 15 ml of TBS buffer (150 mM NaCl; 10 mM Tris pH 7.5) for 15 min on Mixer 820 at speed 60 Hz 18 rpm/8 rpm at RT. Incubation with horseradish peroxidase-conjugated anti-rabbit IgG (1:1000; Sigma-Aldrich, St. Louis, MO, USA) was performed at RT for 3 h in 1% alkali-soluble casein solutions. The membrane was washed with TBS for 15 min on Mixer 820 at speed 60 Hz 18 rpm/8 rpm at RT. The WB was visualized with TBS buffer supplemented with peroxidase substrates (0.002% (w/v) o-dianisidine and 0.03% (v/v) hydrogen peroxide) and incubated in the dark at 37 °C for 30 min. Reaction was stopped by washing the membrane with dH_2_O.

## Results and discussion

icDNA from virus gRNA is one of the proofs that cloned and sequenced viral gRNA sequence is correct and can be regarded as complete. The icDNA construct derived from the RGMoV gRNA sequence, specifically the pJET-RGMoV-cDNAp construct [6]] failed to induce virus infection in oat plants (S1 Fig), indicating the importance of correct 5’ and 3’ ends in the construct. In an attempt to determine the UTRs of the viral genome, both 5’ and 3’ RACE (Rapid Amplification of cDNA Ends) methods were employed. However, only the 3’ UTR was successfully identified, revealing that it was 52 nt longer than previously published sequences [4, 6, 25] and 13 nt longer than the recently published complete genome of RGMoV (GenBank ID MW411579.1) (Fig 1A). Despite efforts to develop the icDNA clone pJET-RGMoV-cDNAp-3end, which included the extended 3’ UTR, this construct also failed to induce local or systemic infection in oat plants (S1 Fig). Subsequently, alternative methods were explored to determine the 5’ UTR. Multiple attempts using RACE and various RT-PCR reaction conditions and primers were unsuccessful. Additionally, the RNA circularization method involving T4 RNA ligase did not yield the desired results. Ultimately, the identification of additional sequences in the 5’ UTR end was achieved through the analysis of RGMoV gRNA RNA-seq HTS data. This information suggests the challenges encountered in characterizing the 5’ and 3’ UTRs of the RGMoV genome and highlights the importance of using multiple approaches, such as RNA-seq, to overcome such difficulties.

**Fig 1.**
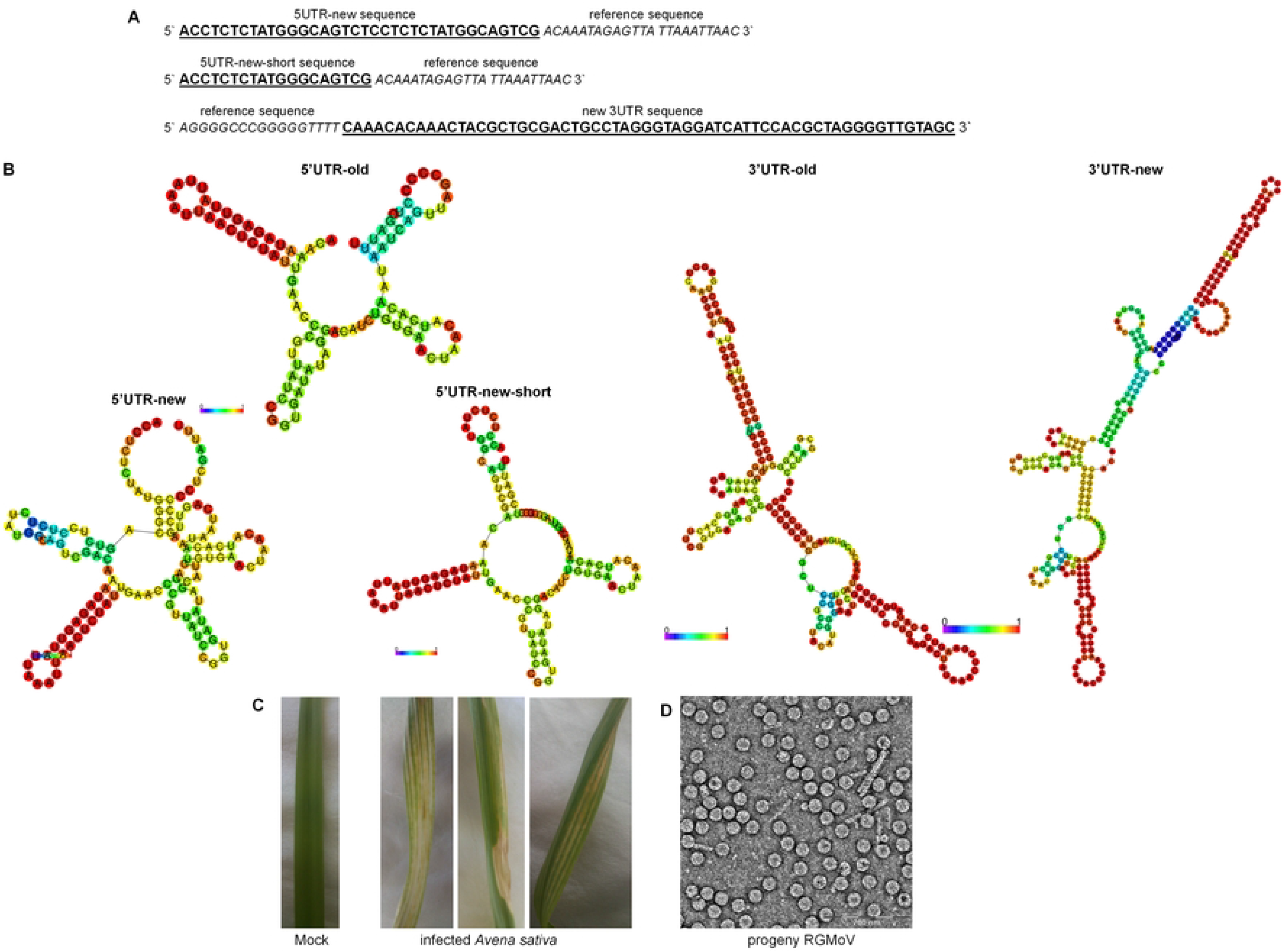
RGMoV icDNA 5′ and 3′ UTR sequence and progeny virus detection. A – schematic sequences of newly identified RGMoV icDNA 5′ and 3′ UTRs; B – comparison of the minimum free energy RNA structures of old and novel identified 5′ and 3′ UTRs created by RNAfold WebServer [39]; C – comparison of Mock and infected *A. sativa* plant leaves bearing RGMoV classical infection pattern; D – transmission electron microscopy analysis of purified progeny virus; bar represents 200 nm scale; reference sequence RGMoV GenBank ID EF091714.1.

Based on the provided information, two sequences were identified for the 5’ UTR of the RGMoV genome: one sequence, referred to as 5UTR-new, was 36 nt longer than the reference sequence (GenBank ID EF091714.1; [6]), and the other sequence, called 5UTR-new-short, was 19 nt longer (Fig 1A). These sequences were identified through manual alignment and analysis of RNA-seq HTS data. It is noted that only one read corresponding to 5UTR-new and two reads corresponding to 5UTR-new-short were identified, with average coverage of 155.9 reads and 176.8 reads, respectively, for the respective UTRs. Attempts were made to construct icDNA clones using these UTR sequences in combination with the 3’ UTR from the reference genome. Specifically, the clones pJET-RGMoV-cDNAp-5UTR-new-PGM and pJET-RGMoV-cDNAp-5UTR-new-short-PGM contained the 5UTR-new and 5UTR-new-short sequences, respectively, along with the reference 3’ UTR. However, both of these constructs failed to induce viral infection (S1 Fig).

Interestingly, the icDNA clone pTZ-T7-RGMoV-cDNA-3end-5UTR-new-short-PGM, which contained the 5UTR-new-short sequence and a new 3’ UTR, was able to induce systemic viral infection with corresponding infection symptoms, such as mottling and necrotic symptoms on leaves (Fig 1C). This outcome is consistent with previous studies [3]. Regarding the 5UTR-new sequence, it is mentioned that it could potentially be an artifact of the HTS library preparation methods or errors during the sequencing process. This suggests the need for further investigation and validation to confirm the authenticity and biological relevance of this sequence.

Overall, these findings highlight the complexities and challenges involved in characterizing UTRs and their impact on viral infectivity, emphasizing the importance of careful experimental design and validation in virus research.

The analysis of possible RNA structures using the RNAfold WebServer [39] has provided valuable insights into the structural features of the 5’ and 3’ UTRs of the RGMoV icDNA. The predicted structure of the 5’ UTR revealed a small loop with a high base-pairing probability (Fig 1B), which may explain the difficulties encountered during 5’ RACE and the low read coverage observed. This small loop structure, in conjunction with the VPg protein, could potentially hinder the efficiency of the 5’ RACE method. On the other hand, the predicted structure of the 3’ UTR of the icDNA exhibited a stem-loop structure with a high base-pairing probability (Fig 1B). This structural feature is absent in all other non-infectious cDNA clones. Similar stem-loop structures have been reported for other sobemoviruses as well [21, 24]. This finding highlights the significance of stem-loop structures in facilitating proper viral genome transcription.

The presence of these distinct structural elements in the UTRs of the RGMoV icDNA suggests their importance in the viral replication cycle and supports their role in facilitating viral genome transcription. Understanding the structural features of the UTRs can provide valuable insights into the mechanisms underlying viral replication and gene expression.

In the study, leaf samples were collected and used for the detection of the coat protein (CP) signal on SDS-PAGE, Western blot (WB) with polyclonal antibodies raised from native virus, and RT-PCR. The results showed that when the gRNA transcript was inoculated by mechanical rubbing, only one out of 10 oat plants showed infection (Fig 2). However, when the gRNA transcript was inoculated using the Helios Gene Gun (HGG) method, six out of 10 oat plants were infected (Fig 2). This demonstrated that the HGG method was more appropriate and effective for *in vitro*-produced viral gRNA inoculation. In the case of an uncapped RNA transcript, no viral infection was detected (Fig 2). This suggests that the presence of a cap structure on the RNA transcript is important for successful viral infection. Furthermore, the study mentioned the development of the first icDNA for sobemovirus, specifically for RYMV. The capped RNA *in vitro* transcript from this icDNA induced a disease phenotype identical to that produced by native viral RNA but the mechanically inoculated RYMV icDNA was found to be less effective. RYMV icDNA RNA transcript without cap showed no disease symptoms four weeks after inoculation [29]. Highlighting the importance of the VPg covalently bound to the RNA in the viral genome transcription process. VPg, found in other plant viruses from the family *Potyviridae*, has been reported as a multifunctional protein involved in viral replication and movement [40]. It interacts with the host cap-binding eIF4E or its isoform (eIF(iso)4E) proteins to initiate viral genome translation [41]. In the case of sobemoviruses, the interaction between RYMV VPg and the central domain of rice eIF(iso)4G1 has been experimentally determined [12, 13]. These findings provide insights into the importance of the viral RNA structure, including the cap structure and VPg, in viral infection and replication processes.

**Fig 2.**
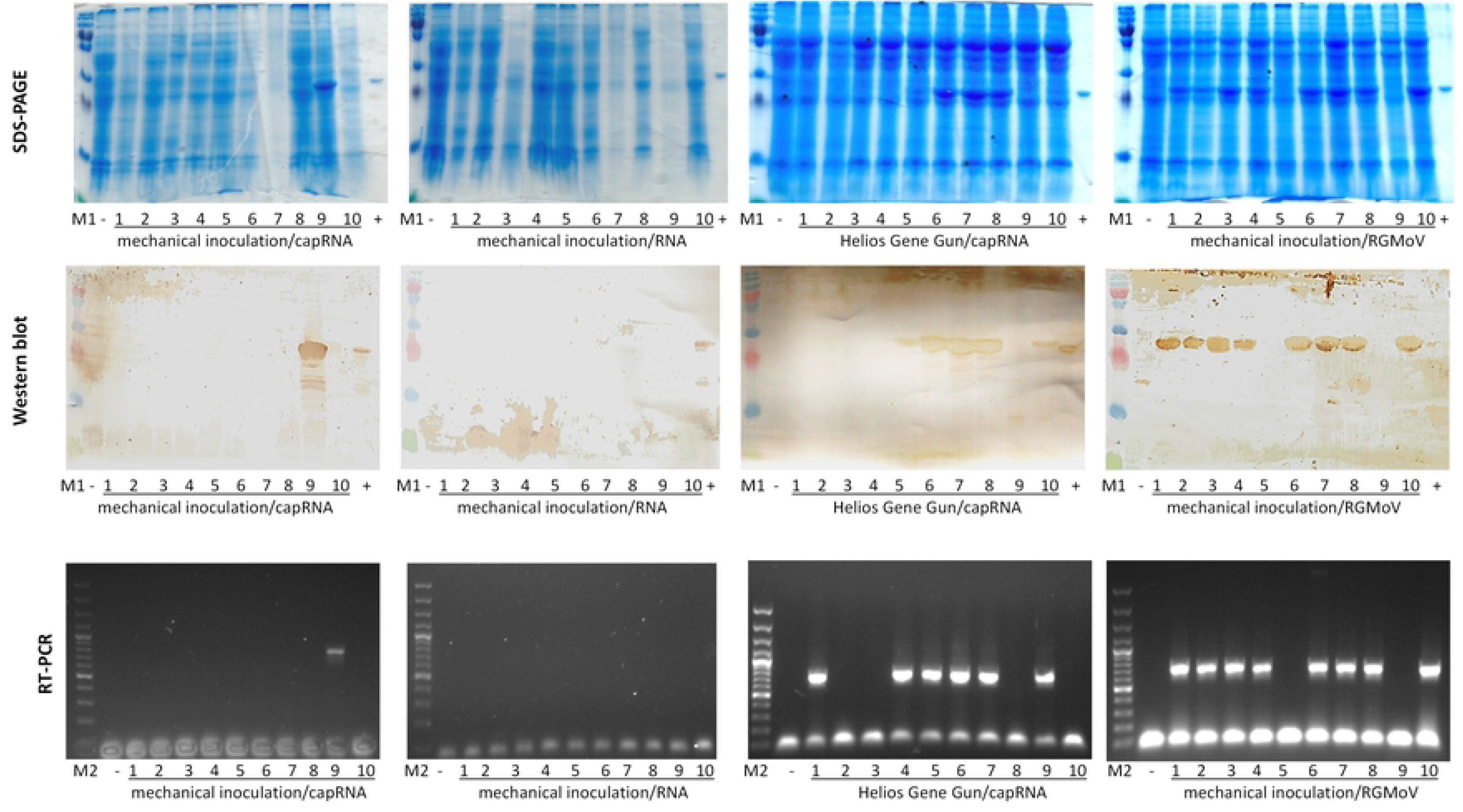
RGMoV icDNA *in vitro* RNA transcript infectivity test in *Avena sativa*. M1 - protein marker (Page Ruler Plus, Thermo Fishers Scientific); M2 – DNA marker (GeneRuler 100 bp Plus DNA Ladder, Thermo Fishers Scientific); “-“ - Mock; 1-10 – inoculated oat plants; “+” - RGMoV as positive control.

TEM analysis of the purified progeny viral fraction revealed typical icosahedral viral particles with an overall diameter of 30 nm (Fig 1D) [5]. This confirms the successful production of intact viral particles. In the study of SeMV, deletion mutants of the 5′ and 3′ UTRs showed that the nucleotide at the 4^th^ position from the 3′ end or the nucleotide at the 5^th^ position from the 5′ end is not crucial for infectivity. However, these mutations resulted in poor accumulation of the CP and unsuccessful purification of progeny virus [16], In the case of RGMoV, CP accumulation was high and progeny virus purification was successful, indicating that the sequenced and cloned infectious cDNA corresponds to the complete viral genome.

The first nucleotide of RGMoV gRNA is a 5′ A, which is similar to RYMV [42] and SBMV [20]. This is in good agreement with the identified VPg binding position at its first serine residue [20]. In the case of RYMV, poorer infectivity was attributed to the presence of an extra 5′ non-viral G residue, which was added to enhance *in vitro* transcription levels [29]. In RGMoV, three additional G residues were added as part of the sequences from position +4 to +8 downstream of the transcription start site, which were found to affect *T7* promoter activity [43]. Similar situations have been observed for other viruses with deviations from the original sequences, such as lacking a native 5′ G residue [44]. In the case of SoMV, the transcribed RNA carried a non-templated G at the 5′ terminus and three non-viral C residues at the 3′ terminus. This RNA was infectious in the test plant *Chenopodium quinoa* but not in *Nicotiana benthamiana* [31]. These findings highlight the importance of specific nucleotide sequences and structures in viral infectivity, CP accumulation, and successful replication. The proper alignment and preservation of the viral genome sequence and structure are crucial for maintaining viral functionality.

The 5′ and 3′ RACE method has been widely used to determine the transcription start point(s) and poly(A) tail sites of mRNA transcripts or viral RNA. It is considered the main strategy for UTR determination [45]. However, there are cases where traditional RACE methods or alternative approaches fail to identify the UTRs accurately. In your study, we found that RNA-seq could be successfully applied to overcome the limitations of traditional methods for UTR determination in the development of icDNA. RNA-seq is a sequencing-based approach that can provide comprehensive information about the transcriptome, including the identification of UTRs. Oxford Nanopore Technologies has described a sequencing approach called 5′ RACE-seq, which combines elements of RACE and RNA-seq to determine the 5′ end of transcripts [45]. Both RACE and RACE-seq methods use adapters to capture and amplify the ends of RNA molecules, allowing for the determination of UTR sequences. The obtained results from these methods have a similar probability of being true, and the accuracy of the results depends on statistical analysis and validation. In our study, the accuracy of the UTR sequences was validated through the propagation of the icDNA and testing its infectivity in progeny virus assays. By utilizing RNA-seq or alternative sequencing approaches, researchers can overcome the limitations of traditional UTR determination methods and obtain more comprehensive and reliable information about the transcriptome, including the UTR sequence.

In our study, we tested the infectivity of progeny virus in oat plants and found that 8 out of 10 infected plants showed evidence of infection through methods such as SDS-PAGE, WB, and RT-PCR (Fig 2). The observed 80% infectivity rate could be attributed to possible genetic variability among the oat seedlings used in the experiment.

To gain a deeper understanding of the observed infectivity and to identify potential differences that may contribute to the infectivity rate, further analysis could be conducted. Transcriptome RNA-seq analysis or analysis of eIF sequences from both infected and non-infected plants could be performed. This analysis could help identify any genetic or transcriptional differences between the plants that may be associated with the infectivity of the progeny virus. It could also shed light on the impact of RGMoV on the transcriptional patterns of host genes, providing valuable insights into the plant’s response to viral infection.

This study opens up new avenues for investigating various aspects of RGMoV, including its replication, RNA packaging, interactions with the host, genome modifications, transmission, symptomatology, and host range. The use of icDNA and the analysis of progeny viruses provide researchers with a powerful tool for studying the genomic and viral life cycle dynamics, as well as their impact on host plants.

The study by Rutgers *et al*. (1980) demonstrated that purified viral particles of southern bean mosaic virus (SBMV) contained minor amounts of sgRNA, which were capable of inducing the synthesis of the CP in wheat embryo extracts and reticulocyte lysates. Similar findings were reported for tymoviruses and tobacco mosaic virus, where minor amounts of sgRNA were encapsidated, while bromoviruses encapsidated sgRNA in significant quantities [46-48]. However, Cho and Dreher showed that sgRNA of the tymovirus turnip yellow mosaic virus was not encapsidated into viral particles [49]. In the case of sobemoviruses, such as LTSV, RYMV, SCMoV, SNMoV, and VTMoV, satRNAs can be encapsidated along with the gRNA. CfMV was found to encapsidate satRNA of LTSV in a host-dependent manner [21], and the encapsulation of defective interfering RNA (diRNA) was also observed [22]. Analyzing HTS data for the identification of encapsulated sgRNA is challenging because the sgRNA sequence is embedded within the gRNA. In the case of RGMoV, the 5’ end of the sgRNA could potentially start at position 2844, where a complementary sequence of five nucleotides from the 5’ UTR (ACAAA) is located. A similar sequence was identified in CfMV, suggesting a hypothetical start of sgRNA [50]. Higher coverage of certain regions in the CP (S2 Fig) coding sequence compared to other regions in the gRNA could indirectly indicate the presence of sgRNA in the analyzed sample, as higher availability of CP matrix sequence suggests the possible presence of sgRNA. Differences in coverage could be attributed to various factors, such as more efficient amplification of certain regions or protection of those regions by CP from RNases during virus storage. These regions may be more abundant and potentially interact with the inner cavity of the virion, potentially serving as origins of assembly (OAS) during virus assembly. As we observed that after virus treatment with RNase A, the extracted RNA concentration was too low for HTS application. OAS or packaging signals were identified in several plant, animal viruses, and phages [51-56]. However, the identification of OAS in RGMoV gRNA in previous experiments was unsuccessful. The siRNA-seq method could be employed for OAS identification.

TEM analysis of RGMoV also revealed the presence of particles with symmetry resembling *T=1* (S3 Fig), suggesting the possibility of encapsulating sgRNA within them. This phenomenon was observed in BMV, where encapsulated RNA controlled the capsid structure. The gRNA was packaged in 180 subunit particles, while mRNA containing only BMV CP was packaged in 120 subunit particles. This demonstrates that RNA features can influence the ribonucleoprotein complex and alternative structural pathways [57]. This could be explained by SBMV sgRNA encapsidate rate (Rutgers, Salerno-Rife et al. 1980). The structural differences in RGMoV CP compared to other sobemoviruses with known CP 3D structures, such as CfMV [58], RYMV [59, 60], SeMV [61], SBMV [62] and Southern cowpea mosaic virus (SCPMV)[60, 63]. may also support the hypothesis of sgRNA encapsidation. Notably, the absence of the FG loop in RGMoV CP decreases the size of the virion cavity by 7% [5].

In conclusion, the encapsidation of sgRNA in viral particles is a complex phenomenon observed in various viruses. Further investigations, including the analysis of RNA-seq data, siRNA-seq, and structural studies, can provide valuable insights into the presence and role of sgRNA in RGMoV and shed light on its impact on viral replication, assembly, and other aspects of the virus life cycle.

That is a significant achievement, as it marks the first report of an infectious and mechanically transmittable full-length cDNA clone of RGMoV. The successful determination of the full genome sequence of RGMoV provides researchers with an important tool for studying sobemoviruses in terms of plant reverse genomics and virus life cycle investigations.

This newly developed methodological approach, combining 5’ RACE-seq and short-read sequencing platforms like Ion PGM, demonstrates the compatibility and efficacy of 5’ RACE-seq in generating comprehensive sequence information. The utilization of this approach enhances our understanding of various aspects of sobemovirus biology, including replication, RNA packaging, host-virus interactions, genome modifications, transmission mechanisms, symptomatology, and host range.

Overall, the availability of the full-length cDNA clone and the complete genome sequence of RGMoV opens up new avenues for further research and provides a valuable resource for the scientific community studying sobemoviruses. Moreover, the cloned virus genome copy paves the way to develop a new vector system for the expression of foreign proteins in plants.

## Acknowledgements

MAFF Genebank is acknowledged for the RGMoV isolate. We thank Dace Skrastiņa from Latvian Biomedical Research and Study Centre, the Biomedical technology complex, the Laboratory animal core facility for polyclonal antibodies of RGMoV.

## Supporting information

**S1 Fig.** Oat leaf sample analysis after inoculation of RGMoV in vitro synthetized gRNA. M1 - protein marker (Page Ruler Plus, Thermo Fishers Scientific); M2 – DNA marker (GeneRuler 100 bp Plus DNA Ladder, Thermo Fishers Scientific); «-» - mock; 1-12 – inoculated oat plants; «+» - RGMoV as positive control.

**S2 Fig.** Schematic read count coverage of RGMoV genome.

**S3 Fig.** RGMoV analysis by TEM. Red arrows indicate possible RGMoV *T=1* particles; white bar represents 500 nm scale.

## Notes

### Competing Interest Statement

The authors have declared no competing interest.

